# Long-term disruption of glucose homeostasis in a rodent model of preterm birth

**DOI:** 10.1101/2024.02.07.579307

**Authors:** Sihao Diao, David Guenoun, Shiou-Ping Chen, Céline Cruciani-Guglielmacci, Julien Pansiot, Mireille Laforge, Ilyes Raho, Valérie Faivre, Vincent Degos, Pierre Gressens, Agnès Nadjar, Juliette Van Steenwinckel, Homa Adle-Biassette, Christophe Magnan, Alice Jacquens, Cindy Bokobza

## Abstract

Around 1 of every 10 babies is born preterm, and the incidence of preterm birth has been rising. The long-term consequences of preterm survivors are not fully understood. Preterm birth is proven to be associated with metabolic diseases and related disorders later in life. Preterm newborns are susceptible to perinatal inflammatory events such as chorioamnionitis, hypoxia-ischemia, and sepsis. We hypothesized that perinatal inflammation has a role in the developmental programming of metabolic diseases and related disorders. In the present study, perinatal inflammation was modeled by systemic administration of IL-1β in mice. We observed a pronounced sexual dimorphism where only the males presented significant insulin resistance and glucose intolerance accompanied by leptin resistance in the long term following perinatal inflammation exposure. Adiposity and energy homeostasis were intact. It showed that perinatal inflammation selectively contributes to the long-term dysregulation of glucose metabolism in a sex-dependent manner. The underlying mechanism might be linked with hypothalamic inflammation and upregulated circulating CCL5. Metformin treatment might be optional to treat insulin resistance resulting from perinatal inflammation.

**Highlights:**

- Perinatal inflammation is common in preterm infants, often leading to perinatal brain injuries. However, the long-term metabolic outcomes of these infants are not fully revealed.
- We explored the long-term metabolic outcomes in mice with perinatal IL-1β exposure and sought its association with inflammation.
- Perinatal inflammation has a profound and deleterious role in glucose metabolism in a sex-dependent and time-dependent manner.
- Perinatal inflammation might be a risk factor for metabolic disorders in preterm survivors.

## Introduction

World Health Organization (WHO) defines obesity as abnormal or excessive fat accumulation that may impair health. Childhood obesity is a major public health problem, WHO estimated in 2016 that 340 million children/adolescents are overweight/obese [1]. Risk factors to develop pediatric obesity are in general associated with food intake by children [2]. However, recent evidence indicates that events during pregnancy or during the first year of infants’ lives also impact the development of childhood obesity [3]. Preterm birth, defined as the livebirth before 37 completed gestational weeks, has become a global health issue. Every year, around 15 million newborns are born preterm, accounting for over 10 % of the total births worldwide [4].With the advance of neonatal medicine and increased survival rate of preterm infants, more and more long-term sequelae have emerged and been drawn attention to in this research field [5, 6]. Previous human cohort studies have indicated that preterm birth predisposes to multiple metabolic diseases and related disorders later in life: former preterm individuals are prone to develop metabolic syndrome, type 2 diabetes mellitus (T2DM), and obesity when compared with peers born at full term [7, 8].

Systemic and hypothalamic inflammation are implicated in the pathogenesis of metabolic disorders and diseases [9, 10]. For example, diet-induced chronic and low-grade inflammation can disrupt insulin signaling in targeted organs (i.e., adipose tissue, hypothalamus, liver, pancreas, muscle) *via* various cytokines and chemokines (TNF-α, IL-1β, IL-6, …) leading to diabetes and related disorders [11]. Inflammation can trigger preterm birth, worsen its outcome, and last long [reviewed in 12, 13]. Based on the above-described evidence, we raise the question of the impact of perinatal inflammation on long-term metabolic vulnerability.

This study used a previously published model of perinatal inflammation associated with preterm birth [14–19]. It is based on intraperitoneal (i.p.) injections of the proinflammatory cytokine IL-1β from postnatal day (P) 1 to P5 to investigate the metabolic outcomes of IL-1β exposure in young and middle-aged animals. In this model, we also evaluated the evolution of inflammation at the periphery and in the central nervous system (CNS). Finally, we tested metformin administration to prevent the metabolic and inflammatory outcomes triggered by IL-1β exposure (Figure (Fig.)1).

## Materials and Methods

### Animals

Experimental protocols were approved by the Ethics committee and the services of the French Ministry in Charge of Higher Education and Research (APAFIS #39290-2022110915453711 v12). Mice were housed under a 12-hour light-dark cycle with free access to water and a chow diet (SAFE 150). Food and water were withdrawn only for glucose and insulin tolerance assays. Experiments were performed on OF1 strain mice (Charles River). Sex was determined at birth and confirmed by abdominal examination at weaning. Both male and female pups were injected i.p. twice a day from P1 to P4 and once a day on P5 with 10 μg/kg IL-1β (Miltenyi, 130-101-684) or phosphate-buffered saline (PBS) 1X alone [14, 17]. Animals were monitored to evaluate general health and measure weight weekly before and biweekly after weaning. We evaluated metabolic and inflammatory changes in young (12 weeks) and middle-aged (28 weeks) animals, respectively.

### Indirect calorimetry

Young and middle-aged mice were individually acclimated in calorimetric cages for one day before the indirect calorimetry analysis (Labmaster, TSE Systems GmbH, Bad Homburg, Germany) and allowed unrestricted access to water and food. The automated measurement of energy expenditure (EE), O_2_ consumption, CO_2_ production, total activity in the x, y, and z dimensions, and food consumption was recorded every 20 minutes for 6 consecutive days. EE was calculated according to the Weir equation. The ANCOVA analysis for EE was analyzed using body weight as a covariant [20]. The predicted EE was calculated based on the regression line equation. The respiratory exchange rate (RER) was estimated by the ratio of CO_2_ production to O_2_ consumption. The MoTil2 IR sensor frames in each calorimetry cage determined total activity. Food consumption was measured by a highly sensitive sensor in each food station. The body composition of lean mass, fat mass, free water, and total water content was analyzed by an EchoMRI (EchoMRI™-100H, Houston, USA) before and after the experiment. Values were corrected by lean body mass for calorimetric analysis.

### Glucose and insulin tolerance assays Glucose tolerance test (GTT)

After a 5 h fast, mice were injected i.p. with 3 g/kg of body weight (30 % glucose). Blood glucose level was measured by a glucose meter (Accu-Check, Roche Applied Science) before and 15, 30, 45, 60, and 90 min after the injection. Blood samples were obtained from the tail vein at 0 and 15 min. The plasma was separated after centrifugation at 4,000 rpm for 10 min. Insulin level was measured by a mouse insulin ELISA kit (Crystal Chem, 90080) according to the manufacturer’s instructions. Homeostasis Model Assessment of insulin resistance (HOMA-IR) and beta cell function (HOMA-β) was calculated by using the formulas:

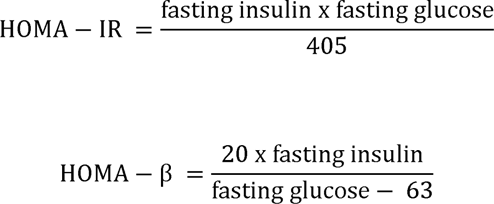

### Insulin tolerance test (ITT)

We proceeded to ITT 48 hours after GTT. After a 5 h fast, mice were injected i.p. with 0.5 IU/kg insulin (NovoRapid FlexPen). Blood glucose level was measured by a glucose meter (Accu-Check, Roche Applied Science) before and 15, 30, 45, 60, and 90 minutes after the injection.

### Luminex assays

Plasma samples were separated by a centrifugation at 1,000 g for 15 minutes at 4 °C and an additional centrifugation at 10,000 for 10 minutes at 4 °C after blood collection from the submandibular vein. Diabetes-related markers and cytokines were measured by Bio-Plex Pro Mouse Diabetes 8-Plex Assay (#171F7001M) and Bio-Plex Pro Mouse Cytokine 23-plex Assay (#M60009RDPD) according to the manufacturer’s instructions (Biorad).

### Flow cytometry

Blood samples were collected as previously described. Immunostaining was performed on 50 uL of blood before hemolysis using BD FACS^TM^ Lysing Solution (BD Biosciences) (activation panel) or on 30 uL of blood after hemolysis (phenotyping panel). The antibodies were listed in supplementary table 1. The samples were kept overnight at 4 °C in the dark, and sample analysis was performed using an LSR Fortessa^TM^ X-20 device (BD Biosciences). We characterized cells as followed: lymphocytes (CD45+ CD11b-), B cells (CD45+ CD11b- CD3- CD45R+), NK cells (CD45+ CD11b- CD3- NK1+), CD4 T cells (CD45+ CD11b- CD3+ CD4+), CD8 T cells (CD45+ CD11b- CD3+ CD8+), neutrophil (SSChi CD11b+ Ly-6g+), monocytes (SSClow CD11b+; Supplementary Fig. 3).

### Immunohistochemistry

Brains were fixed by 4% formaldehyde for 1 week and cut into coronal 16 μm paraffin-embedded sections following dehydration. For immunohistochemistry, the Leica Bond max robot was used with the BOND Polymer Refine Detection kit (Leica, DS9800). Primary antibodies were anti-Iba-1 (Wako, 019-1971, 1:500) and NeuN (Millipore, MAB377, 1:500). For immunofluorescent staining, slices were incubated with blocking solution (0.4 % Triton, 5 % goat serum in PBS) for 1 hour at room temperature after antigen retrieval by heating in citrate buffer. Anti-POMC (Abcam, ab254558, 1:2000) and anti-NeuN (Millipore, MAB377, 1:1000) antibodies were diluted in blocking solution and incubated the slices overnight at 4 °C. Slices were then incubated with secondary antibodies (Cy3 Goat anti-Mouse, Jackson immunoresearch, 115-165-146, 1:1000; Donkey anti-Rabbit, Invitrogen, A-21206, 1:1000) for 1 hour at room temperature, following several washes by PBS. DAPI was used to localize the nuclei of the cells, and then slices were mounted with Fluoromount-G™ Mounting Medium (Invitrogen, 00-4958-02). Immunohistochemistry images were acquired by Nanozoomer slide scanner (Hamamatsu) and analyzed by QuPath [21]. Immunofluorescent images were obtained by a confocal microscope (Leica, TCS SP8) using 40x objectives and analyzed by Fiji with StarDist plugin [22].

### Microglia/macrophage magnetic cell sorting (MACS) and RNA extraction

Animals were perfused intracardially with saline to remove circulating cells. Brains without the cerebellum and olfactory bulbs were collected and dissociated by using an Adult Brain Dissociation Kit and a gentleMACS Octo Dissociator with Heater (Miltenyi Biotec). Magnetic beads coupled with mouse anti-CD11B antibodies (microglia/macrophage) were used for cell isolation according to the manufacturer’s protocol (Miltenyi Biotec) as previously described [14, 23, 24]. According to the manufacturer’s protocol, mRNA from CD11B+ cells was extracted using an RNA XS Plus extraction kit (Macherey-Nagel®).

### RNA sequencing and bioinformatics analysis

Poly(A) RNA sequencing library was prepared following Illumina’s TruSeq-stranded-mRNA sample preparation protocol. Paired-end sequencing was performed on Illumina’s NovaSeq 6000 sequencing system. The analyses were performed using the Eoulsan pipeline, including read filtering, mapping, alignment filtering, read quantification, normalization, and differential analysis as previously described by [18, 23]. StringTie was used to perform mRNA expression levels by calculating FPKM [24]. R package DESeq2 performed mRNAs differential expression analysis between male PBS and IL-1β group [25]. The mRNAs with the parameter of false discovery rate (FDR) below 0.05 and absolute fold change ≥ 2 were considered differentially expressed (DE) mRNAs. Gene Ontology (GO) and Kyoto Encyclopedia of Genes and Genomes (KEGG) pathway enrichment analysis was performed by using the GO database (http://geneontology.org, 2019.05) and KEGG database (http://www.kegg.jp/, 2019.05) respectively. The various gene functional annotations in Fig. 4L were obtained using the Metascape tool v3.5 (http://metascape.org) [26]

### Real-time qPCR

Reverse transcription was performed on 200 ng mRNA using an iScript cDNA synthesis kit (BioRad). Real-time quantitative PCRs were realized by triplicate samples with SYBR Green Supermix (BioRad) as previously described [16, 17]. The primers used in this study were listed in supplementary table 2. mRNA expression levels were calculated by the 2 delta Ct method after normalization with *Rpl13a* as the reference mRNA. Relative expressions were compared with the pre-treatment male PBS group.

### Metformin administration

Middle-aged mice were treated with i.p. 200 mg/kg metformin (Millipore) dissolved in saline for 5 consecutive days. Insulin tolerance tests (ITTs) and flow cytometry were performed, and plasma was collected before and on the 4^th^ day of treatment. CD11B+ cells in the brain were isolated by MACS as described before two hours after the last injection.

### CCL5 ELISA

Platelet-poor plasma was separated immediately after blood collection, following centrifugation at 2,000g for 20 min and an additional centrifugation at 10,000g for 10 min. Mouse CCL5 ELISA Kit (R&D systems, MMR00) was used to quantify CCL5 levels in plasma according to the manufacturer’s protocol.

### Statistical analysis

Data were expressed as mean values with standard deviation (SD). Using GraphPad Prism Software (version 10), single comparisons were made by the Mann–Whitney test. Multiple comparisons in the same data set were analyzed by two-way ANOVA followed by an uncorrected Fisher’s LSD test.

## Results

### Impact of perinatal IL-1β exposure on the body weight and the long-term in vivo metabolism

We evaluated trajectories of the whole-body weight and delta weight gain of the PBS- and IL-1β-exposed animals until middle-age (Fig.2A). We noted a delayed onset of weight gain in male IL-1β-exposed animals from 1 to 4 weeks, and a subsequent significant increase in weight gain from 10 weeks onward compared to PBS-exposed animals. Female IL-1β-exposed animals gained weight later than males, starting at 12 weeks compared to PBS-exposed animals. This difference between males and females is relevant to the clinical features of preterm babies: growth failure caused by perinatal inflammation and the subsequent accelerated “catch-up growth” since adolescence [27]. At young age, we observed that only male IL-1β-exposed animals are overweight compared to PBS-exposed animals (males: p = 0.0032; and females: p = 0.90; Fig.2A). This young age weight gain does not translate in fat over lean mass ratio in both male and female IL-1β-exposed animals compared to PBS-exposed animals (males: P > 0.99; females: P = 0.20; Supplementary Fig.1A). At middle age, we observed that both male and female IL-1β- exposed animals are overweight compared to PBS-exposed animals (males: p = 0.031; and females: p < 0.001; Fig.2C). This middle-age weight gain does not translate to a difference in the fat over lean mass ratio between IL-1β-exposed and PBS-exposed animals in both sexes (males: p = 0.83; females: p = 0.39; Fig.2D). This suggests that perinatal exposure to IL-1β does not affect adiposity in the long term.

To better evaluate global *in vivo* metabolism changes, we placed young and middle-aged PBS- and IL-1β- exposed animals in indirect calorimetry cages for 6 days. In both male and female young and middle-aged mice, we observed no difference in cumulative food intake (FI) in IL-1β-exposed animals compared to PBS-exposed animals (Fig. 2E, Supplementary Fig.1B). In male and female young mice, we observed no difference in total activities in IL-1β-exposed animals compared to PBS-exposed animals (Supplementary Fig.1C/D). In middle-aged male mice, we observed a significant decrease in dark phase IL-1β-exposed animals’ total activities compared to PBS-exposed animals. Such a difference was not observed in females at the same age (Fig. 2F/G).

Total energy expenditure (EE) reflects the sum of basal metabolic rate, thermogenesis from food intake, and energy cost of physical activities [28]. In young male mice, we observed during most of the night an increased EE in IL-1β-exposed animals compared to PBS-exposed animals with a difference between males and females (Figure 2E). To better discriminate between weight-dependent and weight-independent EE, regression-based analysis-of-covariance (ANCOVA) was applied using body weight as a covariant [20]. The deviation of each animal from the regression line is preserved and allows analysis of PBS-exposed and IL-1β-exposed animal differences at a chosen body weight. The ANCOVA analysis of young male mice reveals a weight-independent EE difference with higher EE in the IL-1β-exposed animals compared to PBS-exposed animals (p = 0.0061; Supplementary Fig.1F/H). In male and female middle-aged mice, we observed no difference in EE in IL-1β-exposed animals compared to PBS-exposed animals (males: p = 0.48; females: p = 0.09; Fig.1H-K).

**Figure 1.**
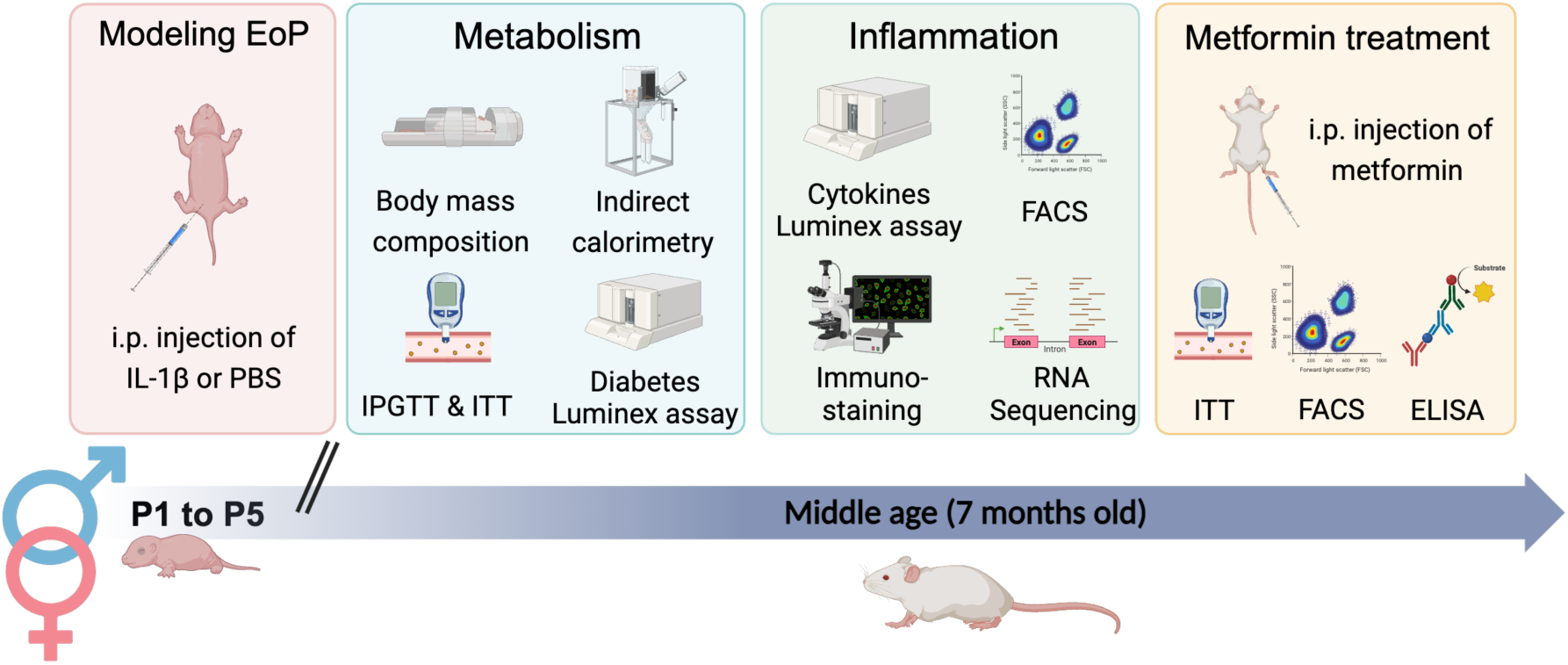
Schematic representation of experimental protocol.

We can extrapolate the Respiratory Exchanges Ratio (RER), a proxy for the animal’s primary fuel source (glycolytic vs. oxidative) from indirect calorimetry. At a young age, we observed no difference in RER in IL-1β-exposed animals compared to PBS-exposed animals (Supplementary Fig.1I/J). However, in middle-aged male mice, we observed a significantly lower RER in IL-1β-exposed animals compared to PBS-exposed animals during the dark period (p = 0.02; Fig.2L/M). Interestingly, this lower RER is simultaneous to lower total activities in male IL-1β-exposed animals (Fig. 2F/L). Overall, we observed excess weight gain and subtle use of lipids as a fuel source in middle-aged male IL-1β-exposed animals.

**Figure 2.**
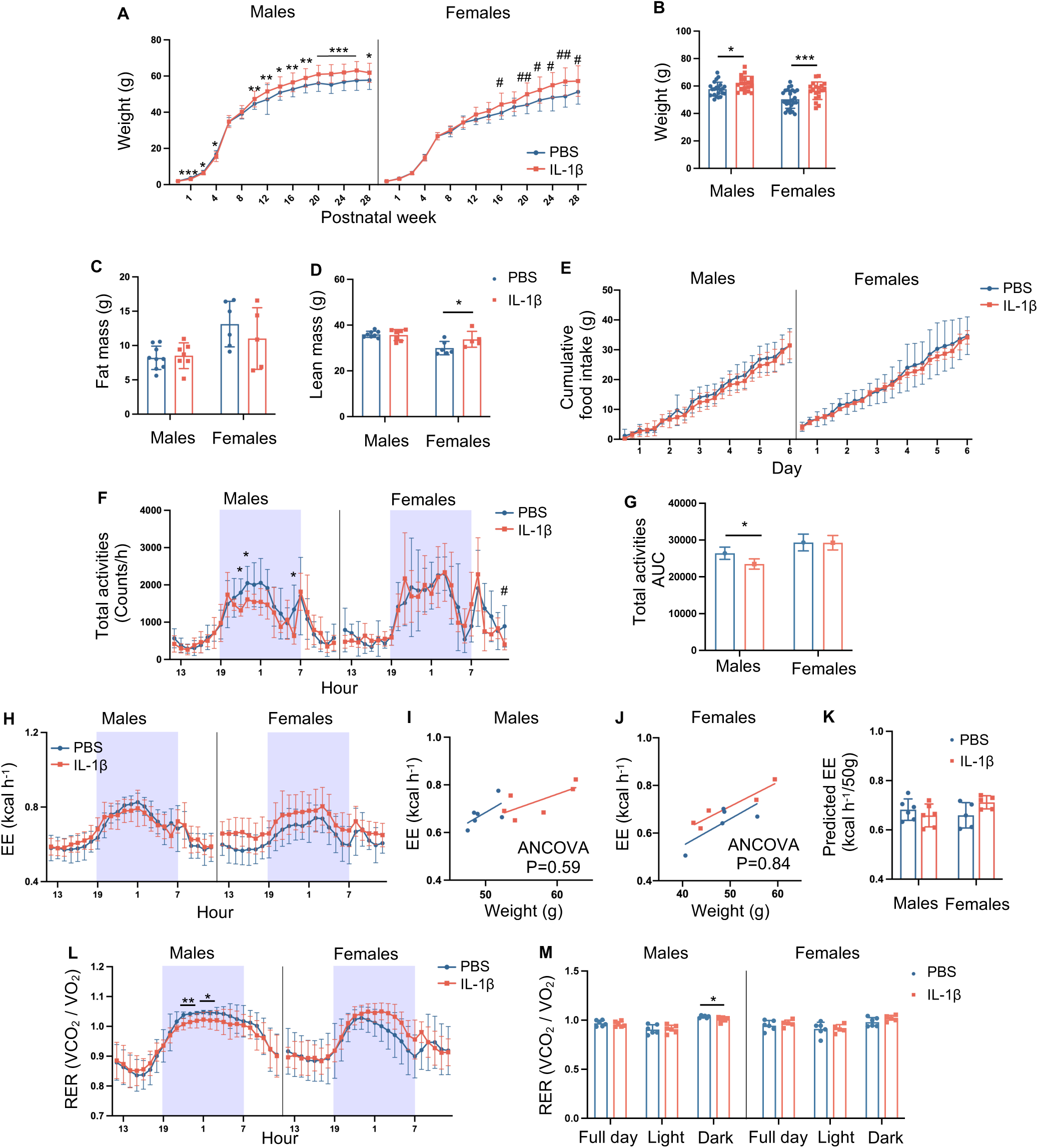
Impact of perinatal IL-1β exposure on the body weight and the long-term in vivo metabolism. (A) Curves of whole-body weight from PBS and IL-1β-exposed mice in males (n = 19 - 42 per group) and females (n = 13 - 32 per group). (B) Whole-body weight of middle-aged PBS- and IL-1β-exposed mice in males (n = 19 - 22 per group) and females (n = 19 - 28 per group). (C) Fat mass of middle-aged PBS- and IL-1β-exposed mice in males (n = 7 - 9 per group) and females (n = 5 - 6 per group). (D) Lean mass of middle-aged PBS- and IL-1β-exposed mice in males (n = 7 - 9 per group) and females (n = 5 - 6 per group). (E) Cumulative food consumption of middle-aged PBS- and IL-1β-exposed mice in males (n = 6 per group) and females (n = 5 per group). (F) Real-time monitoring of total activities of middle-aged PBS- and IL-1β-exposed mice in males (n = 6 per group) and females (n = 5 per group). (G) Quantification of area under the curve of total activities in males and females. (H) Real-time monitoring on EE of middle- aged PBS- and IL-1β-exposed mice in males (n = 6 per group) and females (n = 5 per group). Regression of EE with body weight from middle-aged PBS- and IL-1β-exposed mice in males (I) and females (J). (K) Predicted EE with 50 g body weight for each mouse, calculated by regression lines. (L) Real-time monitoring on RER of middle-aged PBS- and IL-1β-exposed mice in males (n = 6 per group) and females (n = 5 per group). (M) Quantifying RER in full day, light (from 7h to 19h) and dark cycle (from 19h to 7h). Data are presented as mean ± SD. *P* values were determined by 2-way ANOVA followed by Fisher’s LSD test (A-H, K-M) and ANCOVA (I-J). \**P* < 0.05, \*\**P* < 0.01, \*\*\**P* < 0.001, compared with male PBS group. #*P* < 0.05, ##*P* < 0.01, ###*P* < 0.001, compared with female PBS group.

### Impact of perinatal IL-1**β** exposure on the long-term glucose metabolism

Several studies have highlighted a correlation between being born preterm and the disruption of glucose homeostasis at adulthood [8, 29].

We evaluated plasma diabetes-related biomarker levels to explore the effects of perinatal IL-1β exposure on systemic glucose metabolism. At a young age, we observed no difference in evaluated biomarkers in both sexes of IL-1β-exposed animals compared to PBS-exposed animals (Supplementary Fig.2A/B). In middle-aged males, we observed a drastic elevation of leptin levels (p < 0.001) and a significant downregulation of ghrelin (p < 0.001) in the plasma of IL-1β-exposed animals compared to PBS-exposed animals (Fig.3A). Leptin and ghrelin, two hormones exerting opposing effects on EE, play crucial roles in regulating FI. Leptin functions to reduce FI and enhance EE, while ghrelin, an anorexigenic hormone, promotes FI. Despite the absence of discernible disparities in FI and EE among middle-aged males exposed to IL-1β, we posit that leptin and ghrelin may mediate the observed normalization of FI/EE profiles. In middle-aged females, we observed an elevated level of resistin (p=0.008) in the plasma of IL- 1β-exposed animals compared to PBS-exposed animals (Fig.3B). Resitin is an adipocyte, and immune cells secrete a hormone that counteracts insulin effect.

**Figure 3.**
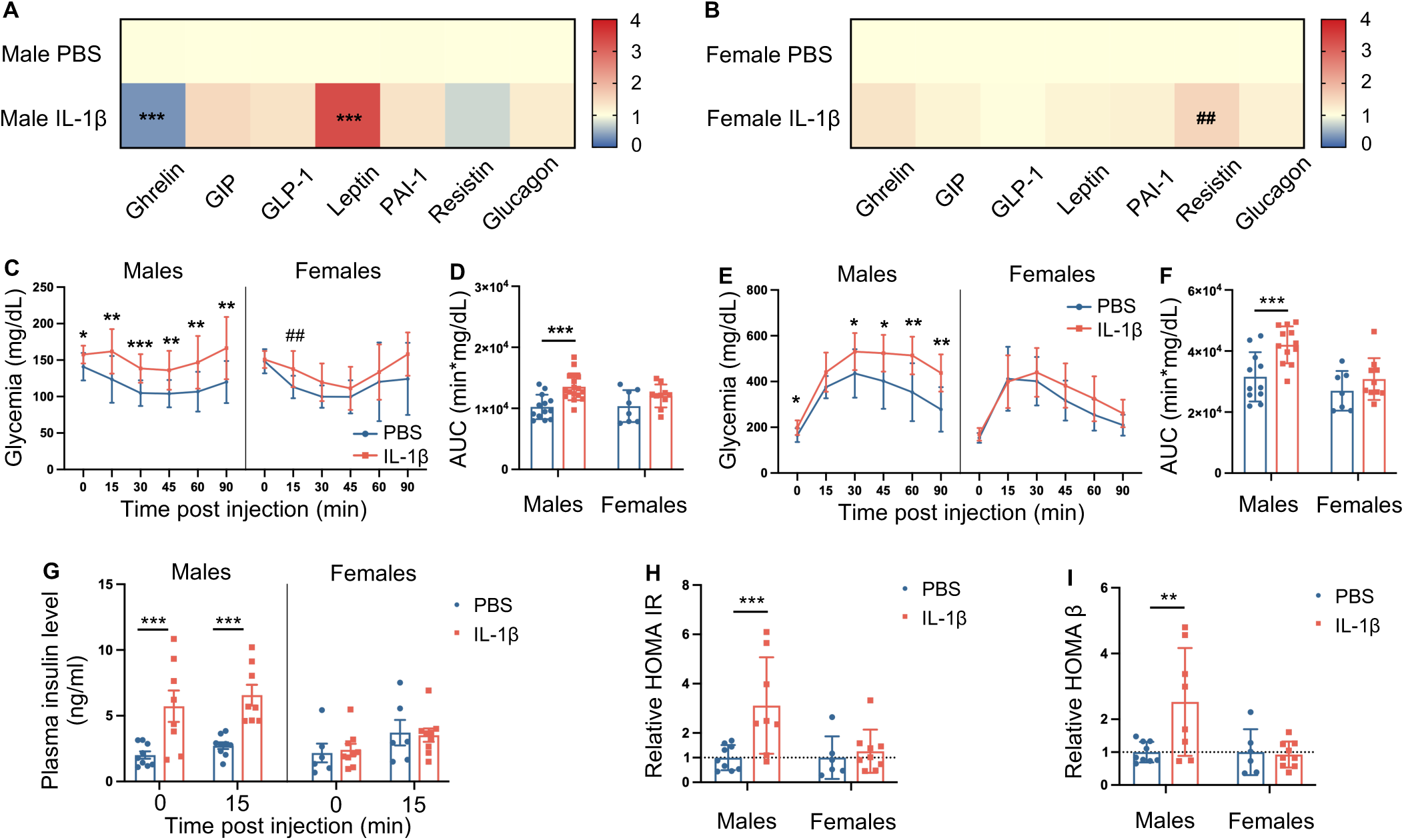
Impact of perinatal IL-1β exposure on the long-term glucose metabolism. Heatmap of relative expression of the diabetes-related markers in plasma of middle-aged PBS- and IL-1β- exposed mice in males (A) (n = 9 - 14 per group) and females (B) (n = 4 - 6 per group). (C) Time course of blood glucose levels during ITT from middle-aged PBS- and IL-1β-exposed mice in males (n = 13 - 17 per group) and females (n = 8 - 10 per group). (D) Quantification of area under the curve of ITT. (E) Time course of blood glucose levels during IPGTT from middle-aged PBS- and IL-1β-exposed mice in males (n = 11 - 12 per group) and females (n = 7 - 10 per group). (F) Quantification of area under the curve of IPGTT. (G) Plasma insulin levels before and 15 min after the injection during IPGTT. Quantification of HOMA-IR index (H) and HOMA-β index (I) normalized to PBS mice in males and females. Data are presented as mean ± SD. *P* values were determined by the Mann-Whitney test (A-B), 2- way ANOVA followed by Fisher’s LSD test (C-I). \**P* < 0.05, \*\**P* < 0.01, \*\*\**P* < 0.001, compared with male PBS group. #*P* < 0.05, ##*P* < 0.01, ###*P* < 0.001, compared with female PBS group.

Hence, we opted to investigate the impact of perinatal IL-1β exposure on insulin sensitivity in young and middle-aged individuals. In young animals, we observed no discernible difference in the response to insulin injection between IL-1β-exposed and PBS-exposed animals in both sexes, even though glycemic normalization appeared swifter in IL-1β-exposed females (Supplementary Fig. 2C/D). In middle-aged males, hyperglycemia manifested before insulin injection after a 5-hour fasting in IL-1β-exposed animals compared to PBS-exposed animals (p = 0.024; Fig. 3C). Additionally, these IL-1β-exposed males displayed significant insulin resistance, as indicated by the absence of glycemic modulation, in stark contrast to PBS-exposed animals (Fig. 3D). Middle-aged females exhibited increased hyperglycemia 15 minutes post-insulin injection in IL-1β-exposed animals as opposed to PBS-exposed counterparts (p = 0.007; Fig. 3C). This transient insulin insensitivity observed in IL-1β-exposed females is consistent with elevated plasmatic resistin levels (Fig. 3B).

Subsequently, we assessed systemic glucose tolerance following perinatal IL-1β exposure in both young and middle-aged individuals. In the younger cohort, no disparities in the response to glucose injection were observed between IL-1β-exposed and PBS-exposed animals in both sexes (Supplementary Fig. 2E/F). In middle-aged males, hyperglycemia was evident after a 5-hour fast in IL-1β-exposed animals compared to PBS-exposed counterparts (p = 0.014; Fig. 3E). Furthermore, IL-1β-exposed males exhibited significant glucose intolerance, as evidenced by the lack of glycemic modulation, in stark contrast to PBS-exposed animals (Fig. 3E/F). There were no discernible differences in the response to glucose injection in middle-aged females between IL-1β- and PBS-exposed animals (Fig. 3E/F). Plasmatic evaluation of insulin levels pre- and post-glucose administration revealed hyperinsulinemia only in middle-aged IL-1β-exposed males (p < 0.001; Fig. 3G, Supplementary Fig. 2G). Fasting glycemia and plasmatic insulin levels were used to calculate HOMA-IR and HOMA-β indexes, respectively assessing insulin resistance and insulin secretion [30]. No differences were observed for both indexes at a young age and in both sexes between IL-1β- and PBS-exposed animals (Supplementary Fig. 2H/I). However, in line with previous findings in middle-aged males only, a significant increase in both HOMA-IR and HOMA-β indexes was observed for IL-1β-exposed animals compared to PBS-exposed animals (p < 0.001 and p < 0.01, respectively; Fig. 3H/I).

In summary, these findings suggest that perinatal IL-1β exposure results in glucose intolerance and insulin resistance coupled with hyperinsulinemia in middle-aged males.

### Impact of perinatal inflammation on the long-term peripheral inflammation and neuroinflammation

The role of inflammation is tightly correlated with the pathogenesis of insulin resistance and glucose intolerance [11]. Perinatal IL-1β exposure is induced by systemic administration and induced at 1 week an elevated plasmatic level of pro-inflammatory cytokines [14, 31] that translated into neuroinflammatory processes mediated by microglia reactivity [32]. We investigated an adult persistent inflammatory syndrome in PBS- and IL-1β-exposed individuals both at the periphery and in the hypothalamus (key brain region regulating glucose metabolism).

We assessed the plasmatic concentrations of various inflammatory-related cytokines and chemokines in middle-aged animals perinatally exposed or not to IL-1β. Among middle-aged females, no discernible differences were observed in the circulating levels of inflammatory molecules between IL-1β- and PBS-exposed animals (Supplementary Fig. 4A). However, among middle-aged males, we noted a slight elevation in the levels of several cytokines and a significant increase in CCL5 in IL-1β-exposed animals compared to PBS-exposed animals (p=0.022; Fig. 4A). To deeper analyze circulating immune cells profiles, we performed blood flow cytometry analysis in adult animals exposed or not to perinatal inflammation. We neither in males nor in females observed any difference between IL-1β- and PBS-exposed animals in percentage of lymphocytes and myeloid circulating cells (Fig. 4B-L, Supplementary Fig. 4B-V). Reactive oxygen species (ROS) are small molecules produced by activated immune cells [33, 34]. Hereby, we evaluated ROS production by neutrophils and monocytes in middle-aged animals. In middle-aged females, we observed no difference in ROS production between PBS- and IL-1β-exposed animals (Supplementary Fig. 4S-V). In middle-aged males, we observed a significant increase of ROS produced by neutrophils and monocytes in IL-1β-exposed animals compared to PBS-exposed animals (p=0.024 and p=0.038, respectively; Fig. 4G-J).

Previous work demonstrated that perinatal IL-1β exposure induced transcriptional changes in microglia/macrophages (CD11B+ cells), brain resident macrophages, that sustain up to 40 days post-IL- 1β administration [15]. Hereby, we magnetically sorted CD11B+ cells from the brains of middle-aged males exposed perinatally to PBS or IL-1β and evaluated transcriptional changes (Fig.4K/L, Supplementary Fig. 4A-C). We observed that 122 genes were differentially expressed by CD11B+ cells (89 downregulated and 33 upregulated) in IL-1β-exposed animals compared to PBS-exposed animals (Fig.4K). The functional annotation using Gene Ontology (GO) highlighted that upregulated genes were associated with the “Immune system process” and “Biological regulation”; downregulated genes were associated with the “Cellular process” and “Homeostatic process.” Interestingly, both up- and downregulated genes were associated with GO of the “Metabolic process” (Fig. 4L). The functional annotation using KEGG pathways for all differentially expressed genes showed altered metabolic pathways such as “Insulin resistance,” “Insulin signaling pathway,” “Regulation of lipolysis in adipocytes,” and “Adipocytokine signaling pathway” (Supplementary Fig. 4B/C). Given that the arcuate nucleus (ARC) plays a pivotal role as a master regulator of systemic glucose metabolism and energy homeostasis within the hypothalamus [35], we proceeded to assess the density of microglial cells (IBA-1+ cells) in this specific region (Fig.4M). In middle-aged males, we observed a significant increase of IBA- 1+ cells in the ARC of IL-1β-exposed animals compared to PBS-exposed animals (p=0.015; Fig. 4N). These data suggest that perinatal IL-1β-exposure induced long-term whole-brain neuroinflammation, including the hypothalamus, is translated by microglial transcriptomic changes coupled to metabolic processes.

**Figure 4.**
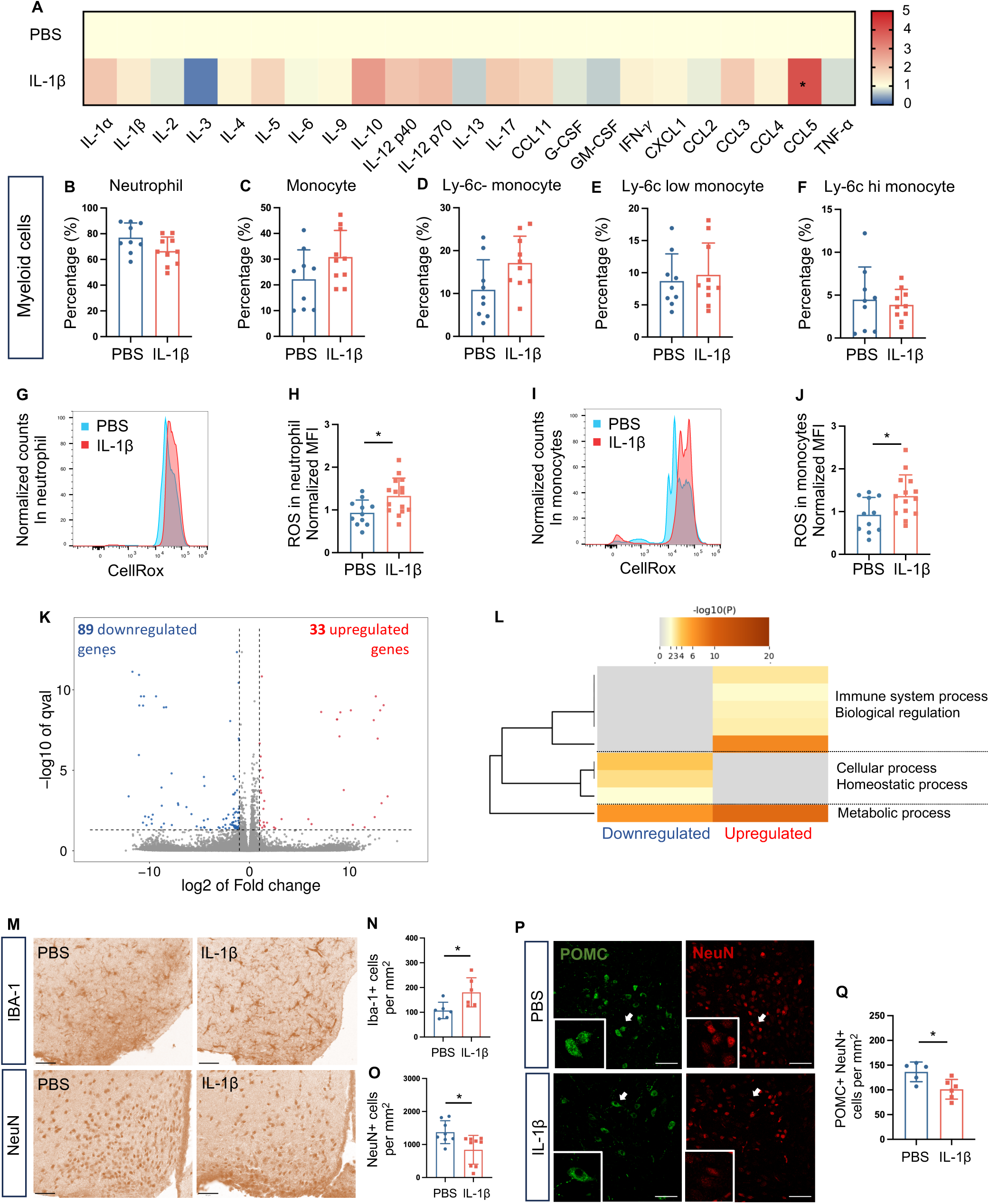
Impact of perinatal inflammation on long-term peripheral inflammation and neuroinflammation. (A) Heatmap of relative expression of cytokines and chemokines in plasma samples of middle-aged PBS- and IL-1β-exposed male mice (n = 9 - 14 per group). Flow cytometry analysis of neutrophils (B), monocytes (C), Ly-6c- monocytes (D), Ly-6c low monocytes (E) and Ly-6c hi monocytes (F) on peripheral blood of middle-aged PBS- and IL-1β-exposed male mice (n = 9 - 10 per group). The fluorescent intensity of ROS in neutrophils (G-H) and monocytes (I-J) of middle-aged PBS- and IL-1β- exposed male mice. (K) Volcano plot illustrating the upregulated and downregulated DEGs of microglia between middle-aged PBS- and IL-1β-exposed male mice. (L) Gene functional annotation of DEGs of microglia between middle-aged PBS- and IL-1β-exposed male mice. (M) Representative immunohistochemistry staining of Iba-1(upper) and NeuN (lower) in the ARC from middle-aged PBS- and IL-1β-exposed male mice (scale bar 50 μm). Quantification of the Iba1+ (N) and NeuN+ (O) cell number per area (n = 6 - 9 per group). (P) Representative immunofluorescence staining of POMC and NeuN in the ARC from middle-aged PBS- and IL-1β-exposed male mice (scale bar 50 μm). (Q) Quantification of POMC+ and NeuN+ cell number per area in the ARC (n = 5 - 6 per group). Data are presented as mean ± SD. *P* values were determined by the Mann-Whitney test. \**P* < 0.05, \*\**P* < 0.01, \*\*\**P* < 0.001.

To further study ARC’s neurons in microgliosis, we performed immunohistochemistry staining of NeuN+ and POMC+ cells in the ARC of middle-aged PBS- and IL-1β-exposed males (Fig. 4O-Q). Our analysis demonstrated a significant decrease in total NeuN+ cells (p=0.011) and POMC+NeuN+ cells (p=0.035) in the ARC of IL-1β-exposed animals compared to PBS-exposed animals, indicating potential disruptions in neuronal populations crucial for glucose homeostasis [36]. These results underscore the intricate relationship between perinatal inflammation, neuroinflammation, and long-term metabolic dysregulations.

### Metformin treatment ameliorated insulin resistance following perinatal inflammation

Metformin is the first-line drug to treat T2DM [37]. It can affect the insulin signaling pathway to improve insulin sensitivity [38]. To explore metformin as a therapeutic strategy for former perinatally inflamed individuals, we tested 5 days treatment (trt) by i.p administration in middle-aged males, in which we observed glucose intolerance (Fig.3). We observed that after 4 days of metformin tx middle-age IL-1β- exposed males do not display any insulin insensitivity (Fig. 5B) and present similar glycemia profiles post-insulin injection to PBS-exposed animals (Fig. 5B). Moreover, we observed a significant reduction in plasma insulin level in fasted middle-age IL-1β-exposed compared to middle-age PBS-exposed males (p = 0.004), suggesting an overall amelioration of insulin resistance (Fig. 4C).

**Figure 5.**
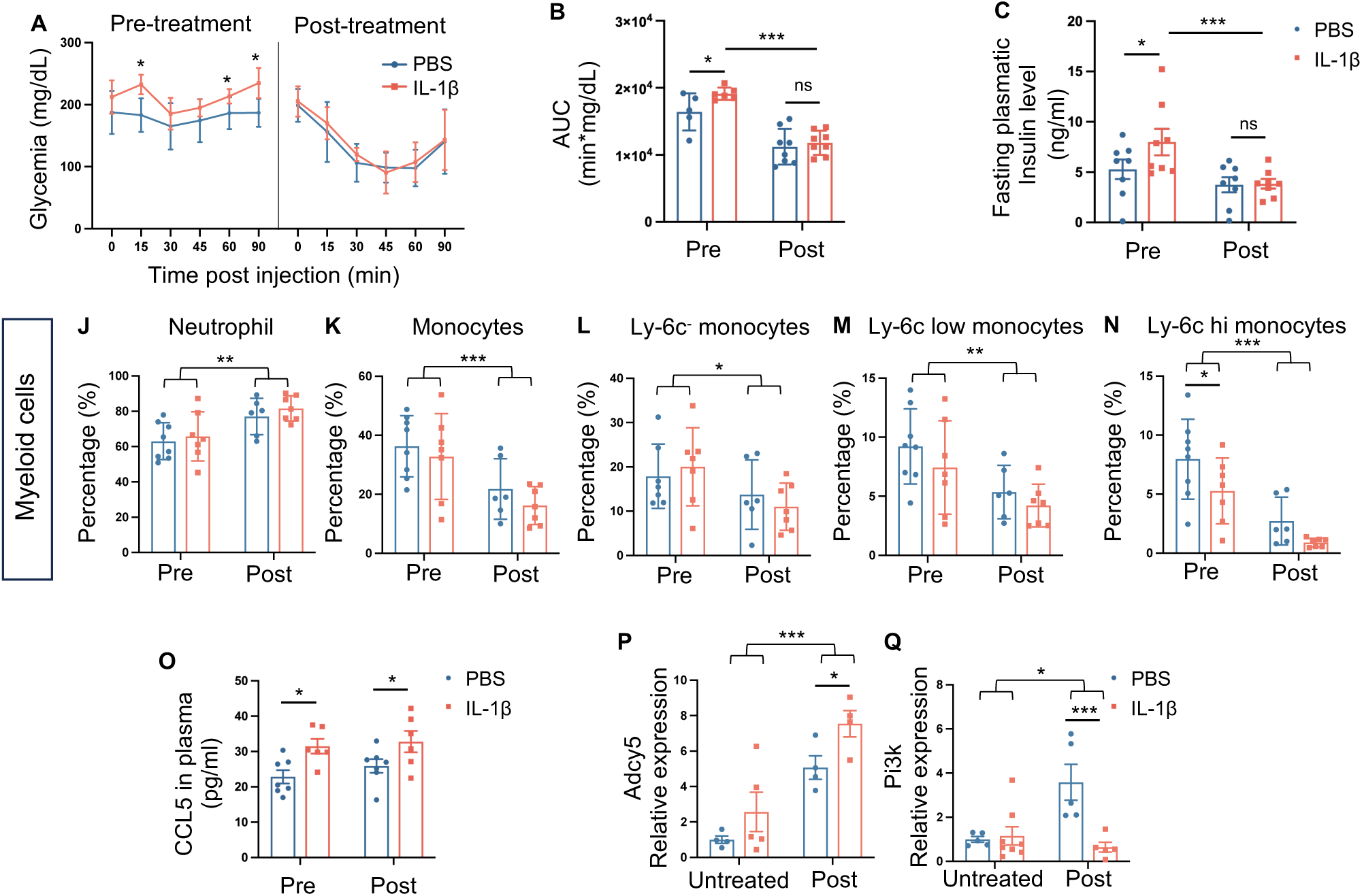
Metformin treatment ameliorated insulin resistance following perinatal inflammation. (A) Time course of blood glucose levels during ITT pre- and post-treatment in middle-aged PBS- and IL- 1β-exposed male mice (n = 5 - 8 per group). (B) Quantification of area under the curve of ITT. (C) Fasting plasma insulin levels pre- and post-treatment in middle-aged PBS- and IL-1β-exposed male mice (n = 8 per group). Flow cytometry analysis of lymphocytes (D), T cells (E), T helper cells (F), cytotoxic cells (G), B cells (H), NK cells (I), neutrophils (J) and monocytes (L-N) on peripheral blood of middle- aged PBS- and IL-1β-exposed male mice pre- and post-treatment. (S) Plasma CCL5 level pre- and post- treatment in middle-aged PBS- and IL-1β-exposed male mice. Relative mRNA expression of Adcy5 (P) and Pi3k (Q) in microglia of middle-aged PBS- and IL-1β-exposed male mice with or without treatment. Data are presented as mean ± SD. P values were determined by Mann-Whitney test (A), 2-way ANOVA followed by Fisher’s LSD test (B – U). \**P* < 0.05, \*\**P* < 0.01, \*\*\**P* < 0.001.

To explore whether metformin trt affects myeloid circulating immune cells, we performed flow cytometry before and on the fifth-day post-trt. We observed a significant reduction of total lymphocyte cells in post-trt IL-1β-exposed males compared to pre-trt IL-1β-exposed males (p=0.038; Supp Fig. 8). We observed no difference in all conditions (Fig. 5E-I). In regards of myeloid cells, we observed a significant increase of neutrophils and reduction of monocytes in post-trt PBS- and IL-1β-exposed males compared to pre-trt PBS- and IL-1β-exposed males (p=0.0013 and p=0.0009 respectively; Fig.5J-N). Interestingly, we observed a significant reduction of Ly-6c^high^ monocytes in pre-trt IL-1β-exposed males compared to pre-trt PBS-exposed animals (p<0.0001) that was counteracted by metformin trt (Fig.5N). Given that CCL5 was the sole circulating cytokine up-regulated in middle-aged IL-1β-exposed males (Fig.4A), we investigated whether metformin treatment could influence its expression levels. Interestingly, in both pre- and post-treatment IL-1β-exposed males, we noted a substantial rise in circulating CCL5 levels compared to PBS-exposed animals, indicating that metformin rectified glucose homeostasis impairments induced by perinatal inflammation without altering the sustained low-grade inflammation linked to CCL5 (Fig.5O).

To explore whether metformin tx affects metabolic signaling pathways altered in microglia/macrophages from IL-1β-exposed males, we evaluated mRNA expression levels of *Adcy5* and *Pi3k*, two genes involved in several of the previously described GO and KEGG pathways link to insulin resistance (Fig.4L, Supplementary Fig. 4B/C)[39]. We observed no difference in *Adcy5* and *Pi3k* mRNA expression levels between untreated PBS- and IL-1β-exposed males. Metformin tx induced overexpression of both *Adcy5* and *Pi3k* mRNA in PBS- and IL-1β-exposed males (p=0.0001 and p=0.043, respectively). In post-tx animals, we observed that CD11B+ cells from IL-1β-exposed males overexpressed and downregulated *Adcy5* and *Pi3k* mRNA expression levels, respectively, when compared to PBS-exposed males (p=0.059 and p=0.0006 respectively, Fig.5P/Q). This suggests that targeting the PI3K pathway might be a good neuroprotective strategy to prevent long-term neuroinflammation.

Taken together, our results demonstrated that metformin treatment might be a strong therapeutic strategy in former inflamed individuals by reducing insulin resistance and some inflammatory parameters that sustain 7 months after perinatal inflammation.

## Discussion

Preterm birth and obesity both represent significant challenges for the World Health Organization (WHO), imposing substantial burdens on clinical, financial, and emotional aspects. In the present study, we evaluated the impact of perinatal inflammation associated with prematurity on the onset of mid- and long-term metabolic dysfunctions using a well-described rodent model.

The primary outcome of our investigation highlights significant sexual dimorphism in long-term metabolism dysfunctions due to perinatal inflammation. We observed a clear “catch-up growth” in male mice, consistent with findings in human cohorts of preterm infants on adiposity, food intake, energy expenditure, and substrate utilization over the long term. However, unlike some human studies reporting increased abdominal adiposity and unhealthy dietary patterns in individuals born preterm [40–42], our study found no differences in adiposity, food intake, and energy expenditure following perinatal IL-1β exposure. We observed enduring dysregulation of glucose metabolism exclusively in male mice following perinatal inflammation, with only subtle impairment observed in their female counterparts. Middle-aged male mice exposed to IL-1β exhibited pronounced insulin resistance and glucose intolerance, resembling metabolic features seen in Type 2 Diabetes Mellitus (T2DM). Additionally, leptin resistance was detected in these middle-aged males exposed to perinatal inflammation. This observation aligns with existing studies indicating that male fetuses/newborns are more susceptible to preterm birth, neonatal morbidity, and poor neurodevelopment outcomes compared to females, possibly linked to sexual differences in placental hormones, brain structural development, and genetics [43–47].

Our work also presents that perinatal inflammation promotes metabolic dysregulation in an age- or time-dependent manner. We observed more subtle changes in young mice than middle-aged mice exposed to perinatal IL-1[. It is consistent with the concept of “Developmental Origins of Health and Diseases,” which proposes the effects of adverse perinatal events in developing metabolic diseases later in life [48]. Previous cohort studies showed that an adverse intrauterine environment impacts glucose homeostasis in a time-dependent manner [49]. The age might reveal the diverse response of the metabolic system to perinatal inflammation stimuli.

Recurrent neuroinflammation was associated with metabolic disorders following perinatal inflammation in the long term. Research has established that perinatal inflammatory events lead to pronounced and diffuse microgliosis in the acute phase [14, 16, 18]. It is noted that our study showed recurrent microgliosis accompanied by neuron loss in the ARC of the hypothalamus and multiple altered metabolic signaling pathways of microglia in the middle-aged IL-1β male group. Mounting evidence suggests that chronic low-grade hypothalamic inflammation, triggered by excessive calorie intake, initiates neuroinflammatory signaling pathways leading to the secretion of pro-inflammatory cytokines and chemokines. This cascade of events contributes to hypothalamic insulin resistance and ultimately fosters the onset of T2DM [50–53]. Our findings suggest a link between hypothalamic inflammation and long-term dysregulation of glucose metabolism resulting from perinatal inflammation. In addition, we noticed a significant upregulation of circulating CCL5 in the middle-aged male IL-1β group. It is proven that patients with T2DM have a higher level of CCL5 [54, 55]. CCL5 is crucial in recruiting immune cells, such as T cells, NK cells, and monocytes, to the inflammatory sites [56]. Recent studies showed that CCL5 knockout mice demonstrated improved glucose tolerance by promoting cell proliferation and insulin secretion under lean conditions [57]. In our study, elevated CCL5 levels might be associated with the dysregulation of glucose metabolism. Further studies would be required to confirm the underlying mechanisms.

Metformin, an FDA-approved drug, has been shown to enhance insulin sensitivity and alleviate low-grade inflammation in individuals with T2DM [58, 59]. In our investigation of the therapeutic potential of metformin in middle-aged IL-1β-exposed male mice, we observed its effective correction of insulin resistance and modulation of circulating myeloid cell populations. Notably, metformin did not impact the IL-1β-induced overexpression of circulating CCL5. These findings suggest that metformin could be beneficial for former preterm adults experiencing glucose homeostasis impairments, offering a promising avenue for managing diabetes. However, it does not directly address the underlying inflammatory mechanisms associated with CCL5, which could perpetuate low-grade inflammation in the periphery or within the hypothalamus. Therefore, further exploration of alternative preventive strategies is necessary to mitigate these inflammatory processes before disruptions in glucose homeostasis occur.

In conclusion, preterm birth and obesity present significant challenges to global health, impacting clinical, financial, and emotional aspects. Our study assessed the influence of perinatal inflammation linked to prematurity on mid- and long-term metabolic dysfunctions using a rodent model. We found notable sexual dimorphism in metabolic dysfunctions due to perinatal inflammation, with male mice exhibiting pronounced insulin resistance and glucose intolerance resembling features of Type 2 Diabetes Mellitus. Recurrent neuroinflammation associated with perinatal inflammation suggests a link between hypothalamic inflammation and long-term dysregulation of glucose metabolism. These findings underscore the need for (i) clinicians to insist on a better follow-up for these specific pediatric populations that are former preterm children and (ii) further research on interventions for long-term metabolic dysfunctions, with metformin emerging as one potential therapeutic option.

## Author Contributions

S.D., D.G., P.G., A.N., J.VS., H.A., A.J. and C.B.: study concept and design. S.D., D.G., S.C., C.C., C.M., J.P., M.L., Z.C., I.S., I.R., V.F., H.A. and C.B.: data acquisition and analysis. S.D., D.G., A.J. and C.B.: drafting manuscript.

## Acknowledgment

Graphical abstract and schematic representations were created using the BioRender tool. We thank Zsolt Csaba, Irvin Sautet, Jennifer Hua, Ariane Heydari-Olya, and the indirect calorimetry platform from Université Paris Cité BFA, for their technical assistance.

## Figures Legends

**Supplementary Figure 1. Impact of perinatal IL-1β exposure on body weight at young age.**

(A) Curves of weight gain of PBS- and IL-1β-exposed mice from in males (n = 19 - 42 per group) and females (n = 13 - 32 per group). (B) Curves of whole-body weight of PBS- and IL-1β-exposed mice from in males (n = 19 - 42 per group) and females (n = 13 - 32 per group) from postnatal week 0 to 4. (C) Curves of weight gain of PBS- and IL-1β-exposed mice from in males (n = 19 - 42 per group) and females (n = 13 - 32 per group) from postnatal week 0 to 4. Data are presented as mean ± SD. *P* values were determined by 2-way ANOVA followed by Fisher’s LSD test. \**P* < 0.05, \*\**P* < 0.01, \*\*\**P* < 0.001, compared with male PBS group. #*P* < 0.05, ##*P* < 0.01, ###*P* < 0.001, compared with female PBS group.

**Supplementary Figure 2. Impact of perinatal IL-1β exposure on body composition and in vivo metabolism at young age.**

(A) Fat mass of young PBS- and IL-1β-exposed mice in males (n = 4 - 8 per group) and females (n = 6 per group). (D) Lean mass of young PBS- and IL-1β-exposed mice in males (n = 4 - 8 per group) and females (n = 6 per group). (C) Cumulative food consumption of young PBS- and IL-1β-exposed mice in males (n = 4 - 8 per group) and females (n = 6 per group). (D) Real-time monitoring of total activities of young PBS- and IL-1β-exposed mice in males (n = 4 - 8 per group) and females (n = 6 per group). (E) Quantification of area under the curve of total activities in males and females. (F) Real-time monitoring on EE of young PBS- and IL-1β-exposed mice in males (n = 4 - 8 per group) and females (n = 6 per group). Regression of EE with body weight from young PBS- and IL-1β-exposed mice in males (G) and females (H). (I) Predicted EE with 38 g body weight for each male mouse or with 28 g body weight for each female mouse, calculated by regression lines. (J) Real-time monitoring on RER of young PBS- and IL-1β-exposed mice in males (n = 4 - 8 per group) and females (n = 6 per group). (K) Quantification of RER in full day, light (from 7h to 19h) and dark cycle (from 19h to 7h). Data are presented as mean ± SD. P values were determined by 2-way ANOVA followed by Fisher’s LSD test (A-F, I-K), ANCOVA (G-H). \**P* < 0.05, \*\**P* < 0.01, \*\*\**P* < 0.001, compared with male PBS group. #*P* < 0.05, ##*P* < 0.01, ###*P* < 0.001, compared with female PBS group.

**Supplementary Figure 3. Impact of perinatal IL-1**β **exposure on glucose metabolism at young age.** Heatmap of relative expression of the diabetes-related markers in plasma of young PBS- and IL-1β- exposed mice in males (A) (n = 3 – 4 per group) and females (B) (n = 3 – 4 per group). (C) Time course of blood glucose levels during ITT from young PBS- and IL-1β-exposed mice in males (n = 21 per group) and females (n = 13 per group). (D) Quantification of area under the curve of ITT. (E) Time course of blood glucose levels during IPGTT from young PBS- and IL-1β-exposed mice in males (n = 13 per group) and females (n = 12 mice per group). (F) Quantification of area under the curve of IPGTT. (G) Plasma insulin levels before and 15 min after the injection during IPGTT. Quantification of HOMA-IR index (H) and HOMA-β index (I) normalized to PBS-exposed mice in males and females. Data are presented as mean ± SD. *P* values were determined by the Mann-Whitney test (A-B), 2-way ANOVA followed by Fisher’s LSD test (C-I). \**P* < 0.05, \*\**P* < 0.01, \*\*\**P* < 0.001, compared with male PBS group. #*P* < 0.05, ##*P* < 0.01, ###*P* < 0.001, compared with female PBS group.

**Supplementary Figure 4. Gating strategy for flow cytometry analysis on the blood**

For the phenotype panel, all cells were first included (A). Doublets were excluded based on SSC (B) and FSC (C). Live cells were gated based on the viability dye (D). The total lymphocyte population was gated based on CD45+ and CD11b- expression (E). T lymphocytes were gated as CD3+ and NK1- population (F). Cytotoxic T cells and T helper cells were gated as CD8+ CD4- and CD4+ CD8- population, respectively (G). B cells and NK cells were gated as CD45R+ NK1- and NK1+ CD45R- population, respectively (H). For the activation panel, all cells were first included (I). Doublets were excluded based on SSC (J) and FSC (K). Live cells were gated based on the viability dye (L). Neutrophils were gated as CD11b+, SSC hi, I-A I-E- and Ly-6G+ cells (M-P). Monocytes were gated as CD11b+, SSC low cells (M-N) and then classified into Ly-6c-, Ly-6c low, and Ly-6c hi monocytes (R). The fluorescent intensity of ROS in neutrophils (Q) and monocytes (S) were demonstrated respectively.

**Supplementary Figure 5. RNA-seq analysis of microglia in middle-aged male mice following perinatal IL-1β exposure.**

(A) Heatmap representation of DEGs from RNA-seq analysis of microglia between middle-aged PBS- and IL-1β-exposed male mice. (B) KEGG enriched analysis of DEGs from RNA-seq analysis of microglia between middle-aged PBS- and IL-1β-exposed male mice. (C) Gene functional annotation of DEGs of microglia between middle-aged PBS- and IL-1β-exposed male mice

**Supplementary Figure 4. Impact of perinatal IL-1β exposure on peripheral inflammation at middle age.**

(A) Heatmap of relative expression of cytokines and chemokines in plasma samples of middle-aged PBS- and IL-1β-exposed female mice (n = 5 - 6 per group). Flow cytometry analysis of lymphocytes (B), T cells (C), CD4+ T cells (D), CD8+ cells (E) and B cells (F) on peripheral blood of middle-aged PBS- and IL-1β-exposed male mice (n = 12 - 17 per group). Flow cytometry analysis of lymphocytes (H), T cells (I), CD4+ T cells (J), CD8+ cells (K), B cells (L), NK cells (M), neutrophils (N) and monocytes (O-R) on peripheral blood of middle-aged PBS- and IL-1β-exposed female mice (n = 9 - 12 per group). The fluorescent intensity of ROS in neutrophils (S-T) and monocytes (U-V) of middle-aged PBS- and IL-1β- exposed female mice (n = 9 - 10 per group). Data are presented as mean ± SD. *P* values were determined by the Mann-Whitney test. #*P* < 0.05, ##*P* < 0.01, ###*P* < 0.001.

**Supplementary Figure 5. Impact of perinatal IL-1β exposure on peripheral inflammation at young age.**

Flow cytometry analysis of lymphocytes (A), T cells (B), CD4+ T cells (C), CD8+ T cells (D), B cells (E), NK cells (F), neutrophils (G) and monocytes (H-K) on peripheral blood of young PBS- and IL-1β-exposed mice in males and females (n = 4 - 6 per group). The fluorescent intensity of ROS in neutrophils (L-N) and monocytes (O-R) of young PBS- and IL-1β-exposed mice (n = 4 - 6 per group). Data are presented as mean ± SD. *P* values were determined by 2-way ANOVA followed by Fisher’s LSD test. \**P* < 0.05, \*\**P* < 0.01, \*\*\**P* < 0.001, compared with male PBS group. #*P* < 0.05, ##*P* < 0.01, ###*P* < 0.001, compared with female PBS group.

**Supplementary Figure 6. Impact of metformin treatment on lymphocytes.**

Flow cytometry analysis of lymphocytes (A), T cells (B), CD4+ T cells (C), CD8+ T cells (D), B cells (E), NK cells (F) on peripheral blood of middle-aged PBS- and IL-1β-exposed male mice pre- and post- treatment. Data are presented as mean ± SD. P values were determined by 2-way ANOVA followed by Fisher’s LSD test. \**P* < 0.05, \*\**P* < 0.01, \*\*\**P* < 0.001.

## References

1. WHO. https://www.who.int/news-room/fact-sheets/detail/obesity-and-overweight. 2019 [cited.

2. Haqq, A.M., et al., Complexity and Stigma of Pediatric Obesity. Child Obes, 2021. 17(4): p. 229–240.

3. Seneviratne, S.N. and S. Rajindrajith, Fetal programming of obesity and type 2 diabetes. World J Diabetes, 2022. 13(7): p. 482–497.

4. Chawanpaiboon, S., et al., Global, regional, and national estimates of levels of preterm birth in 2014: a systematic review and modelling analysis. Lancet Glob Health, 2019. 7(1): p. e37–e46.

5. Marlow, N., et al., Neurologic and developmental disability at six years of age after extremely preterm birth. N Engl J Med, 2005. 352(1): p. 9–19.

6. Nosarti, C., et al., Preterm birth and psychiatric disorders in young adult life. Arch Gen Psychiatry, 2012. 69(6): p. E1–8.

7. Crump, C., J. Sundquist, and K. Sundquist, Preterm birth and risk of type 1 and type 2 diabetes: a national cohort study. Diabetologia, 2020. 63(3): p. 508–518.

8. Markopoulou, P., et al., Preterm Birth as a Risk Factor for Metabolic Syndrome and Cardiovascular Disease in Adult Life: A Systematic Review and Meta-Analysis. J Pediatr, 2019. 210: p. 69–80.e5.

9. Hotamisligil, G.S., Inflammation and metabolic disorders. Nature, 2006. 444(7121): p. 860–7.

10. Bhusal, A., M.H. Rahman, and K. Suk, Hypothalamic inflammation in metabolic disorders and aging. Cell Mol Life Sci, 2021. 79(1): p. 32.

11. Rohm, T.V., et al., Inflammation in obesity, diabetes, and related disorders. Immunity, 2022. 55(1): p. 31–55.

12. Humberg, A., et al., Preterm birth and sustained inflammation: consequences for the neonate. Semin Immunopathol, 2020. 42(4): p. 451–468.

13. Dammann, O. and A. Leviton, Intermittent or sustained systemic inflammation and the preterm brain. Pediatr Res, 2014. 75(3): p. 376–80.

14. Favrais, G., et al., Systemic inflammation disrupts the developmental program of white matter. Ann Neurol, 2011. 70(4): p. 550–65.

15. Krishnan, M.L., et al., Integrative genomics of microglia implicates DLG4 (PSD95) in the white matter development of preterm infants. Nat Commun, 2017. 8(1): p. 428.

16. Van Steenwinckel, J., et al., Decreased microglial Wnt/β-catenin signalling drives microglial pro-inflammatory activation in the developing brain. Brain, 2019. 142(12): p. 3806–3833.

17. Bokobza, C., et al., miR-146b Protects the Perinatal Brain against Microglia-Induced Hypomyelination. Ann Neurol, 2022. 91(1): p. 48–65.

18. Klein, L., et al., A unique cerebellar pattern of microglia activation in a mouse model of encephalopathy of prematurity. Glia, 2022. 70(9): p. 1699–1719.

19. Bokobza, C., et al., Targeting the brain 5-HT7 receptor to prevent hypomyelination in a rodent model of perinatal white matter injuries. J Neural Transm (Vienna), 2023. 130(3): p. 281–297.

20. Muller, T.D., M. Klingenspor, and M.H. Tschop, Revisiting energy expenditure: how to correct mouse metabolic rate for body mass. Nat Metab, 2021. 3(9): p. 1134–1136.

21. Bankhead, P., et al., QuPath: Open source software for digital pathology image analysis. Sci Rep, 2017. 7(1): p. 16878.

22. Schmidt, U., et al. Cell Detection with Star-Convex Polygons. in Medical Image Computing and Computer Assisted Intervention – MICCAI 2018. 2018. Cham: Springer International Publishing.

23. Jourdren, L., et al., Eoulsan: a cloud computing-based framework facilitating high throughput sequencing analyses. Bioinformatics, 2012. 28(11): p. 1542–1543.

24. Pertea, M., et al., StringTie enables improved reconstruction of a transcriptome from RNA-seq reads. Nat Biotechnol, 2015. 33(3): p. 290–5.

25. Love, M.I., W. Huber, and S. Anders, Moderated estimation of fold change and dispersion for RNA-seq data with DESeq2. Genome Biol, 2014. 15(12): p. 550.

26. Zhou, Y., et al., Metascape provides a biologist-oriented resource for the analysis of systems-level datasets. Nat Commun, 2019. 10(1): p. 1523.

27. Embleton, N.D., et al., Catch-up growth and metabolic outcomes in adolescents born preterm. Arch Dis Child, 2016. 101(11): p. 1026–1031.

28. Donahoo, W.T., J.A. Levine, and E.L. Melanson, Variability in energy expenditure and its components. Curr Opin Clin Nutr Metab Care, 2004. 7(6): p. 599–605.

29. Wang, G., et al., Preterm birth and random plasma insulin levels at birth and in early childhood. Jama, 2014. 311(6): p. 587–96.

30. Matthews, D.R., et al., Homeostasis model assessment: insulin resistance and beta-cell function from fasting plasma glucose and insulin concentrations in man. Diabetologia, 1985. 28(7): p. 412–9.

31. Bokobza, C., et al., Targeting the brain 5-HT7 receptor to prevent hypomyelination in a rodent model of perinatal white matter injuries. J Neural Transm (Vienna), 2022.

32. Van Steenwinckel, J., et al., Decreased microglial Wnt/beta-catenin signalling drives microglial pro-inflammatory activation in the developing brain. Brain, 2019. 142(12): p. 3806–3833.

33. Martinvalet, D. and M. Walch, Editorial: The Role of Reactive Oxygen Species in Protective Immunity. Front Immunol, 2021. 12: p. 832946.

34. Liemburg-Apers, D.C., et al., Interactions between mitochondrial reactive oxygen species and cellular glucose metabolism. Arch Toxicol, 2015. 89(8): p. 1209–26.

35. Jais, A. and J.C. Brüning, Arcuate Nucleus-Dependent Regulation of Metabolism- Pathways to Obesity and Diabetes Mellitus. Endocr Rev, 2022. 43(2): p. 314–328.

36. Varela, L. and T.L. Horvath, Leptin and insulin pathways in POMC and AgRP neurons that modulate energy balance and glucose homeostasis. EMBO Rep, 2012. 13(12): p. 1079–86.

37. Sanchez-Rangel, E. and S.E. Inzucchi, Metformin: clinical use in type 2 diabetes. Diabetologia, 2017. 60(9): p. 1586–1593.

38. Herman, R., et al., Metformin and Insulin Resistance: A Review of the Underlying Mechanisms behind Changes in GLUT4-Mediated Glucose Transport. Int J Mol Sci, 2022. 23(3).

39. Gao, P., et al., Underlying Mechanism of Insulin Resistance: A Bioinformatics Analysis Based on Validated Related-Genes from Public Disease Databases. Med Sci Monit, 2020. 26: p. e924334.

40. Sipola-Leppänen, M., et al., Resting energy expenditure in young adults born preterm--the Helsinki study of very low birth weight adults. PLoS One, 2011. 6(3): p. e17700.

41. Sharafi, M., et al., Dietary behaviors of adults born prematurely may explain future risk for cardiovascular disease. Appetite, 2016. 99: p. 157–167.

42. Vohr, B.R., et al., Extreme Preterm Infant Rates of Overweight and Obesity at School Age in the SUPPORT Neuroimaging and Neurodevelopmental Outcomes Cohort. J Pediatr, 2018. 200: p. 132–139.e3.

43. Kelly, L.A., et al., Sex differences in neonatal brain injury and inflammation. Front Immunol, 2023. 14: p. 1243364.

44. Peelen, M.J., et al., Impact of fetal gender on the risk of preterm birth, a national cohort study. Acta Obstet Gynecol Scand, 2016. 95(9): p. 1034–41.

45. Vacher, C.M., et al., Placental endocrine function shapes cerebellar development and social behavior. Nat Neurosci, 2021. 24(10): p. 1392–1401.

46. Kozhemiako, N., et al., Sex differences in brain connectivity and male vulnerability in very preterm children. Hum Brain Mapp, 2020. 41(2): p. 388–400.

47. Benavides, A., et al., Sex-specific alterations in preterm brain. Pediatr Res, 2019. 85(1): p. 55–62.

48. Barker, D.J. and C. Osmond, Infant mortality, childhood nutrition, and ischaemic heart disease in England and Wales. Lancet, 1986. 1(8489): p. 1077–81.

49. Poulsen, P. and A. Vaag, The intrauterine environment as reflected by birth size and twin and zygosity status influences insulin action and intracellular glucose metabolism in an age- or time-dependent manner. Diabetes, 2006. 55(6): p. 1819–25.

50. Arruda, A.P., et al., Low-Grade Hypothalamic Inflammation Leads to Defective Thermogenesis, Insulin Resistance, and Impaired Insulin Secretion. Endocrinology, 2011. 152(4): p. 1314–1326.

51. Ono, H., Molecular Mechanisms of Hypothalamic Insulin Resistance. Int J Mol Sci, 2019. 20(6).

52. Sewaybricker, L.E., et al., The Significance of Hypothalamic Inflammation and Gliosis for the Pathogenesis of Obesity in Humans. Endocr Rev, 2023. 44(2): p. 281–296.

53. Leon, S., A. Nadjar, and C. Quarta, Microglia-Neuron Crosstalk in Obesity: Melodious Interaction or Kiss of Death? Int J Mol Sci, 2021. 22(10).

54. Dworacka, M., et al., Increased circulating RANTES in type 2 diabetes. Eur Cytokine Netw, 2014. 25(3): p. 46–51.

55. Pan, X., et al., Chemokines in Prediabetes and Type 2 Diabetes: A Meta-Analysis. Front Immunol, 2021. 12: p. 622438.

56. Yao, M., et al., Chapter Eight - Cytokine Regulation of Metastasis and Tumorigenicity, in Advances in Cancer Research, D.R. Welch and P.B. Fisher, Editors. 2016, Academic Press. p. 265–367.

57. Zhou, H., et al., Metabolic effects of CCL5 deficiency in lean and obese mice. Front Immunol, 2022. 13: p. 1059687.

58. He, L., Metformin and Systemic Metabolism. Trends Pharmacol Sci, 2020. 41(11): p. 868–881.

59. Feng, X., et al., Metformin, Macrophage Dysfunction and Atherosclerosis. Front Immunol, 2021. 12: p. 682853.

