## Supplementary Table for "Long-term disruption of glucose homeostasis in a rodent model of preterm birth"

**Supplementary Table 1. List of antibodies for flow cytometry.**

| **Phenotype panel** | | | | |
| --- | --- | --- | --- | --- |
| Target | Conjugate | Clone | Company | Catalog No. |
| CD45 | VioBlue | REA737 | Miltenyi | 130-110-802 |
| CD11b | VioBright FITC | REA592 | Miltenyi | 130-113-805 |
| CD3 | APC Vio770 | REA606 | Miltenyi | 130-117-788 |
| NK1 | PE | REA1162 | Miltenyi | 130-120-506 |
| CD4 | BV785 | RM4-5 | Biolegend | 100552 |
| CD8a | PE Vio 615 | REA601 | Miltenyi | 130-123-914 |
| CD45R | PerCP Vio 700 | REA755 | Miltenyi | 130-110-850 |
| **Activation panel** | | | | |
| Target | Conjugate | Clone | Company | Reference No. |
| I-A I-E | BV711 | M5/114.15.2 | Sony | 1138215 |
| Ly-6G | PE Cy7 | 1A8 | Biolegend | 127617 |
| CD11b | BV421 | M1/70 | Sony | 1106255 |
| Ly-6C | BV605 | HK1.4 | Biolegend | 128036 |

**Supplementary Table 2. List of primers used in this study.**

| Primers | Sequences 5’ to 3’ | Products (bp) |
| --- | --- | --- |
| Adcy5_Fw | TGAACCAGAGCAGTCTCACCAT | 75 |
| Adcy5_Rv | GCGCCGCGTGGAAAG | 75 |
| Pi3k_Fw | TACTTGATGTGGCTGACGCA | 122 |
| Pi3l_Rv | TCATGGTGGGGCAAATCCTC | 122 |
