## Supplementary figures for "Long-term disruption of glucose homeostasis in a rodent model of preterm birth"

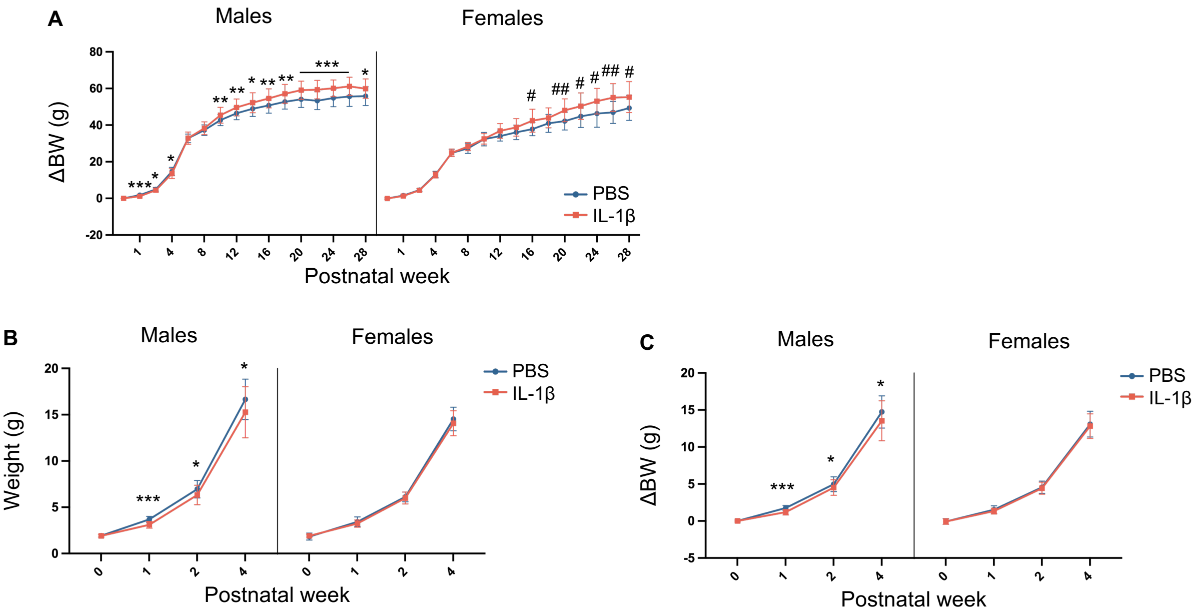
A
Males
Females
ΔBW (g)
80
60
40
20
0
-20
1
4
8
12
16
20
24
28
***
*
**
#
##
Postnatal week
PBS
IL-1β
B
Males
Females
Weight (g)
20
15
10
5
0
0
1
2
4
***
*
Postnatal week
PBS
IL-1β
C
Males
Females
ΔBW (g)
20
15
10
5
0
-5
0
1
2
4
***
*
Postnatal week
PBS
IL-1β

### Supplementary Figure 2

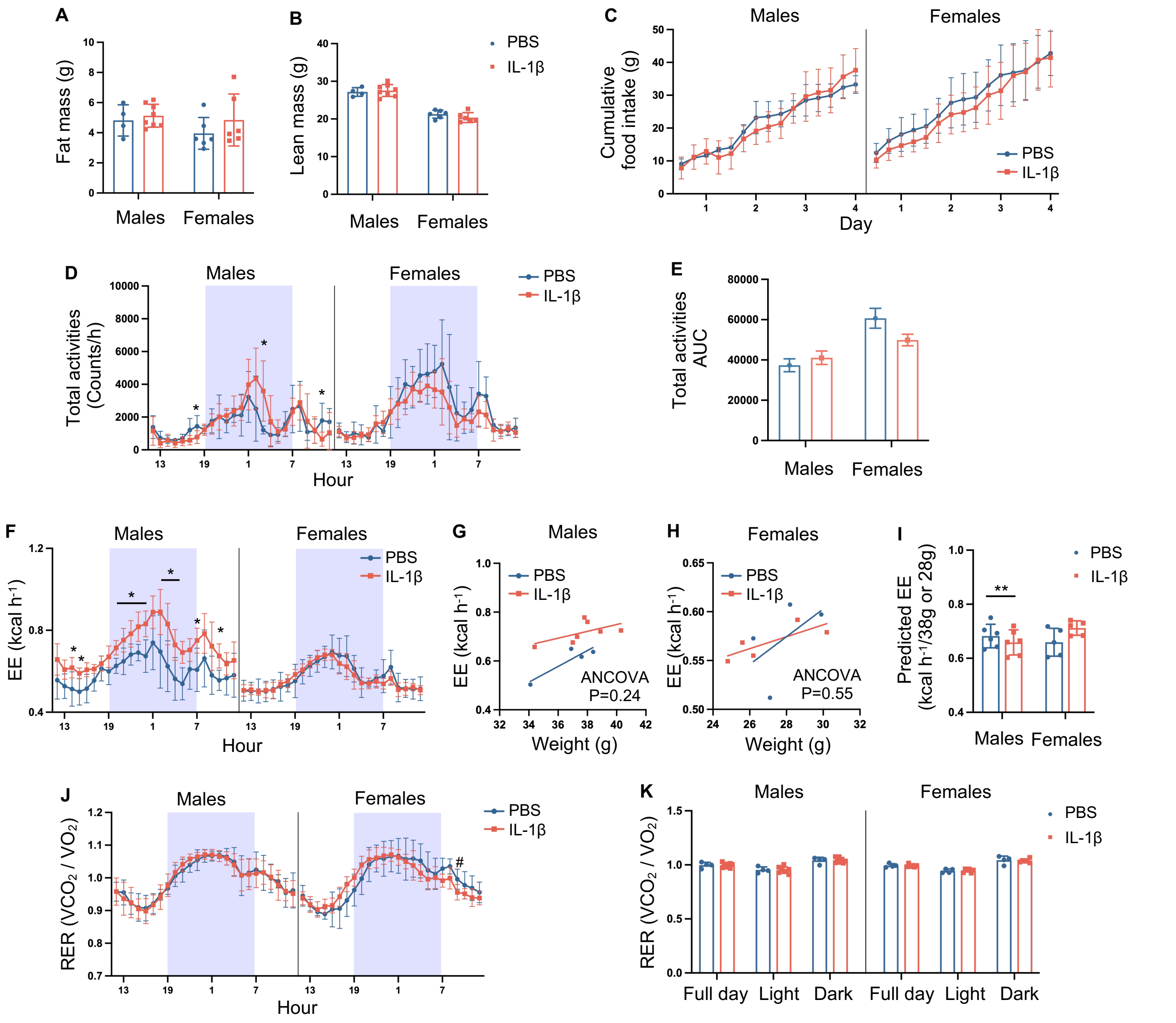

### Supplementary Figure 3

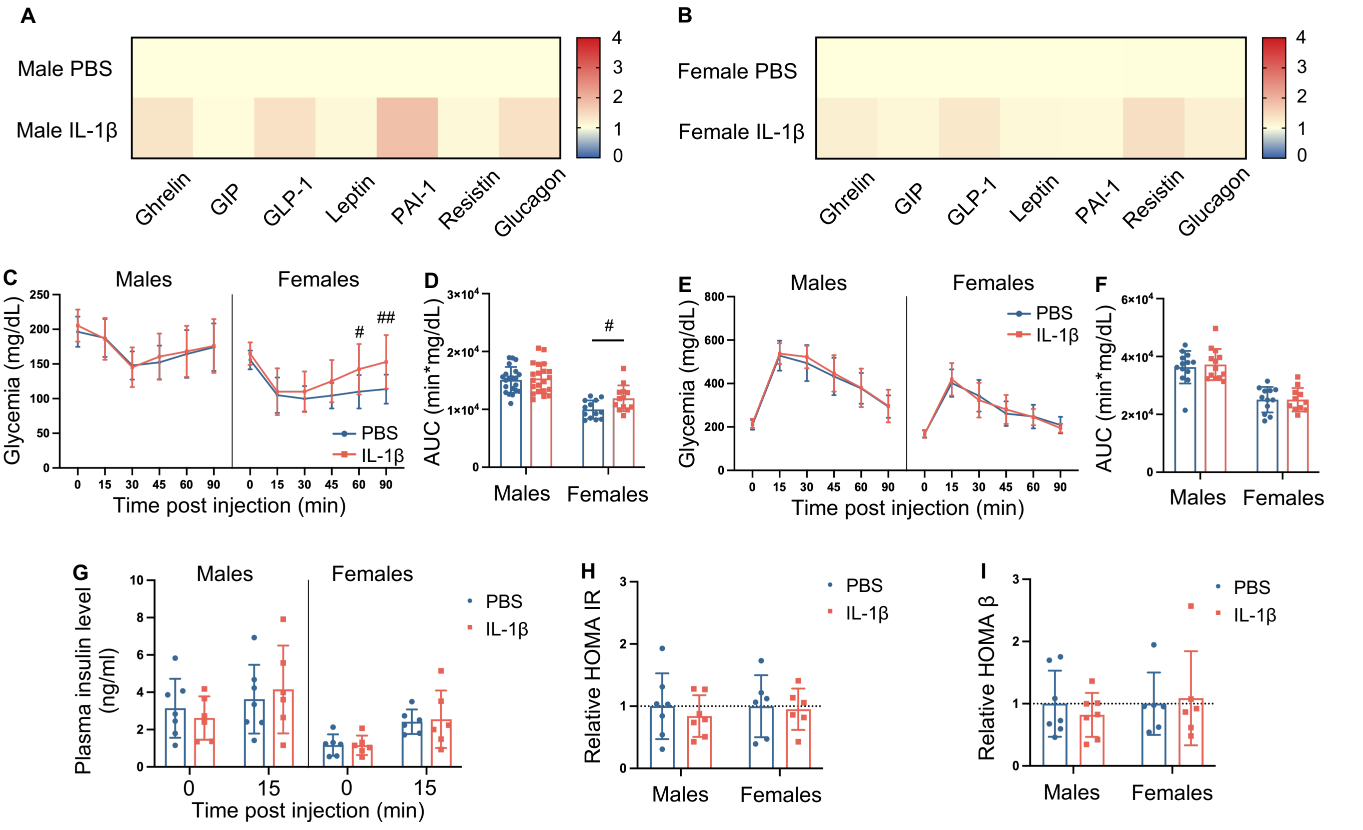

### Supplementary Figure 4

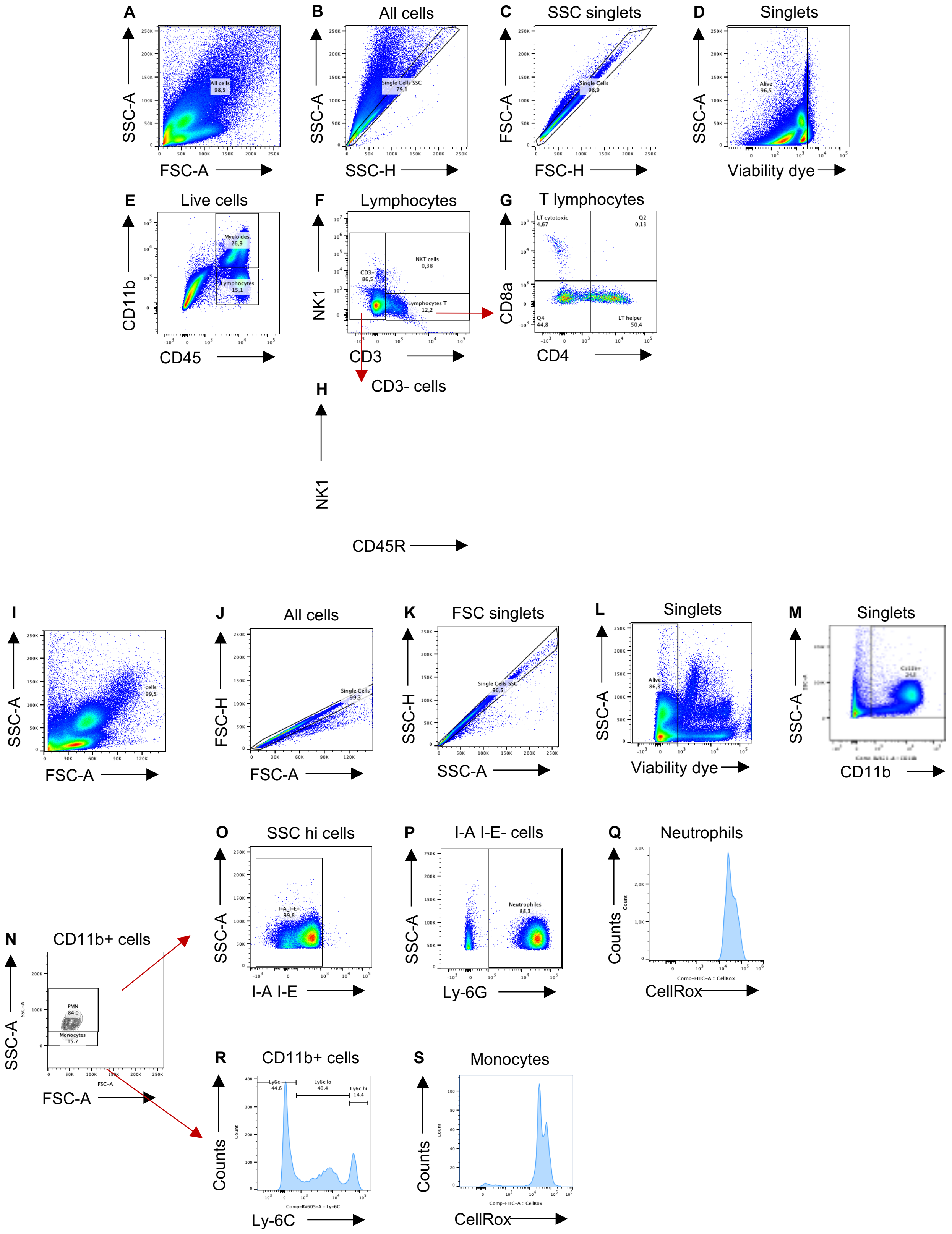

### Supplementary Figure 5

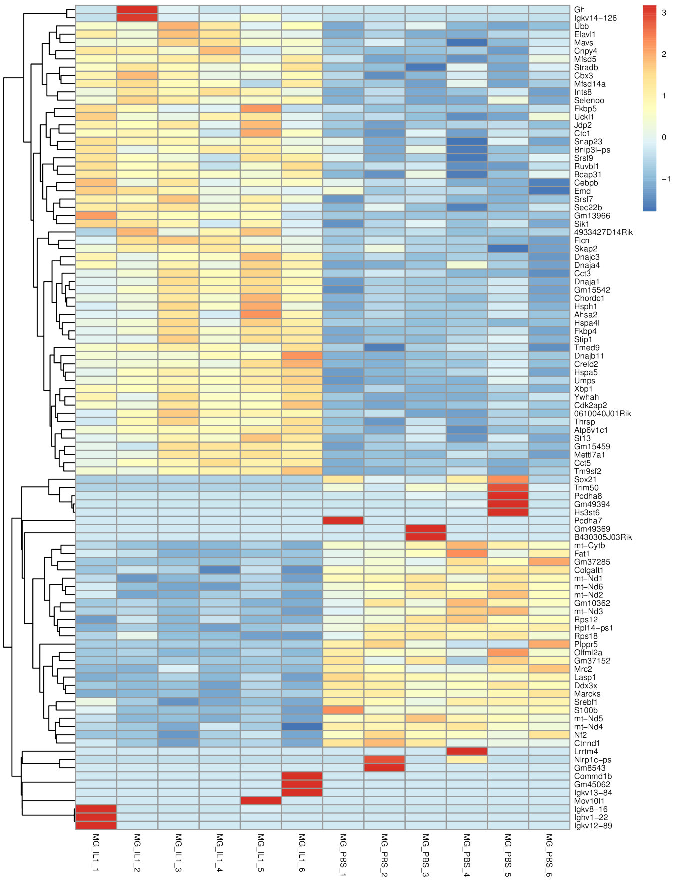


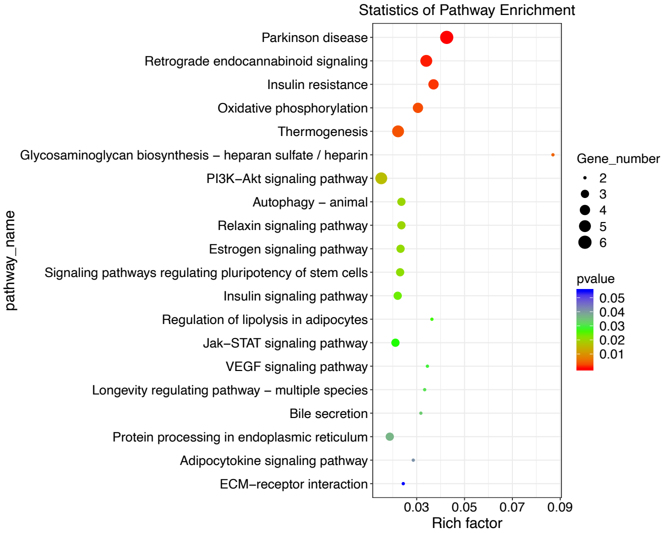


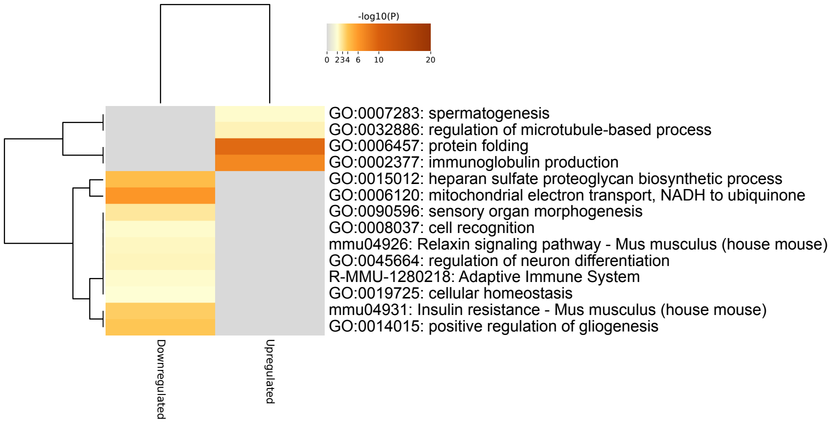

### Supplementary Figure 6

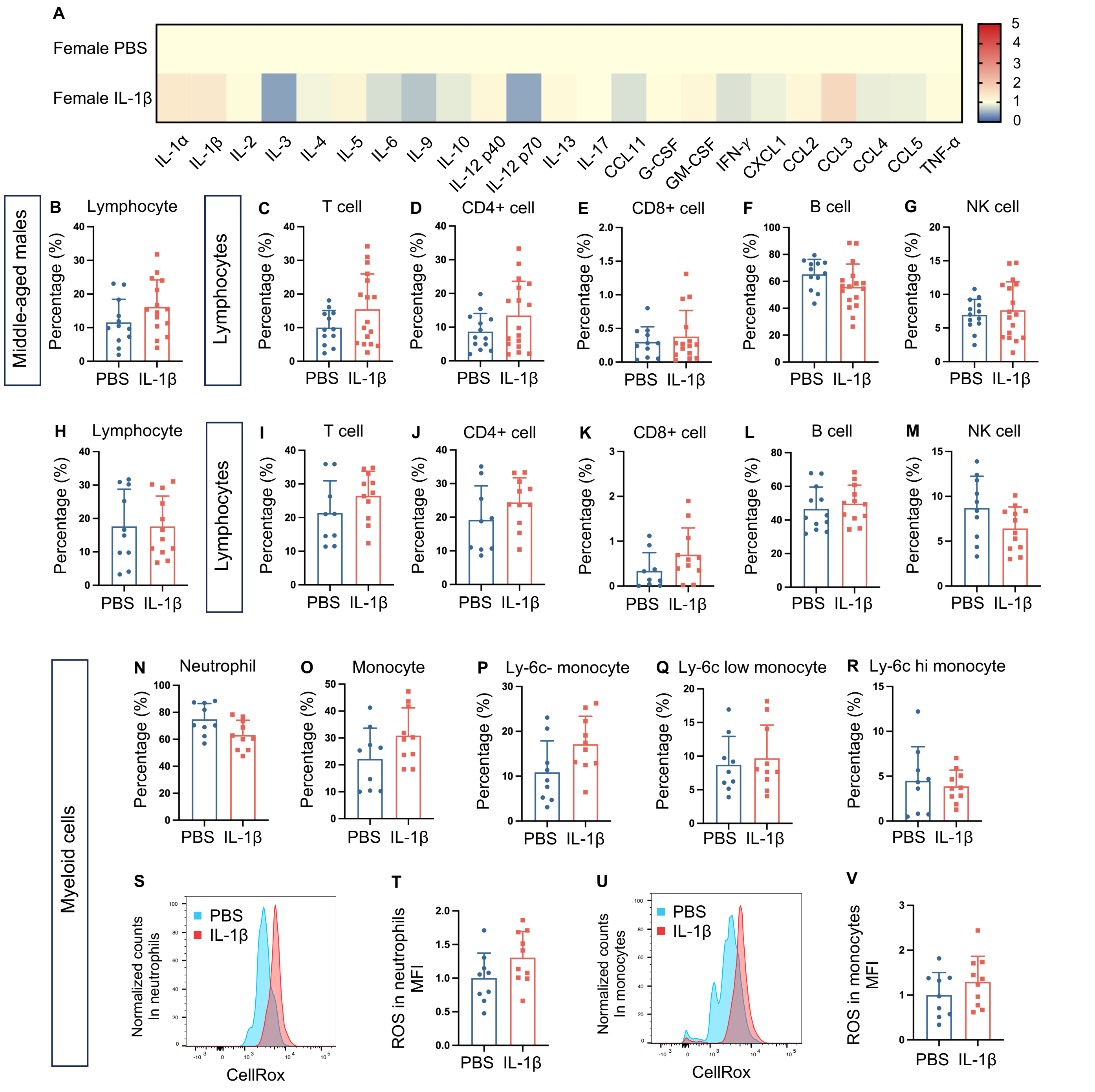

### Supplementary Figure 7

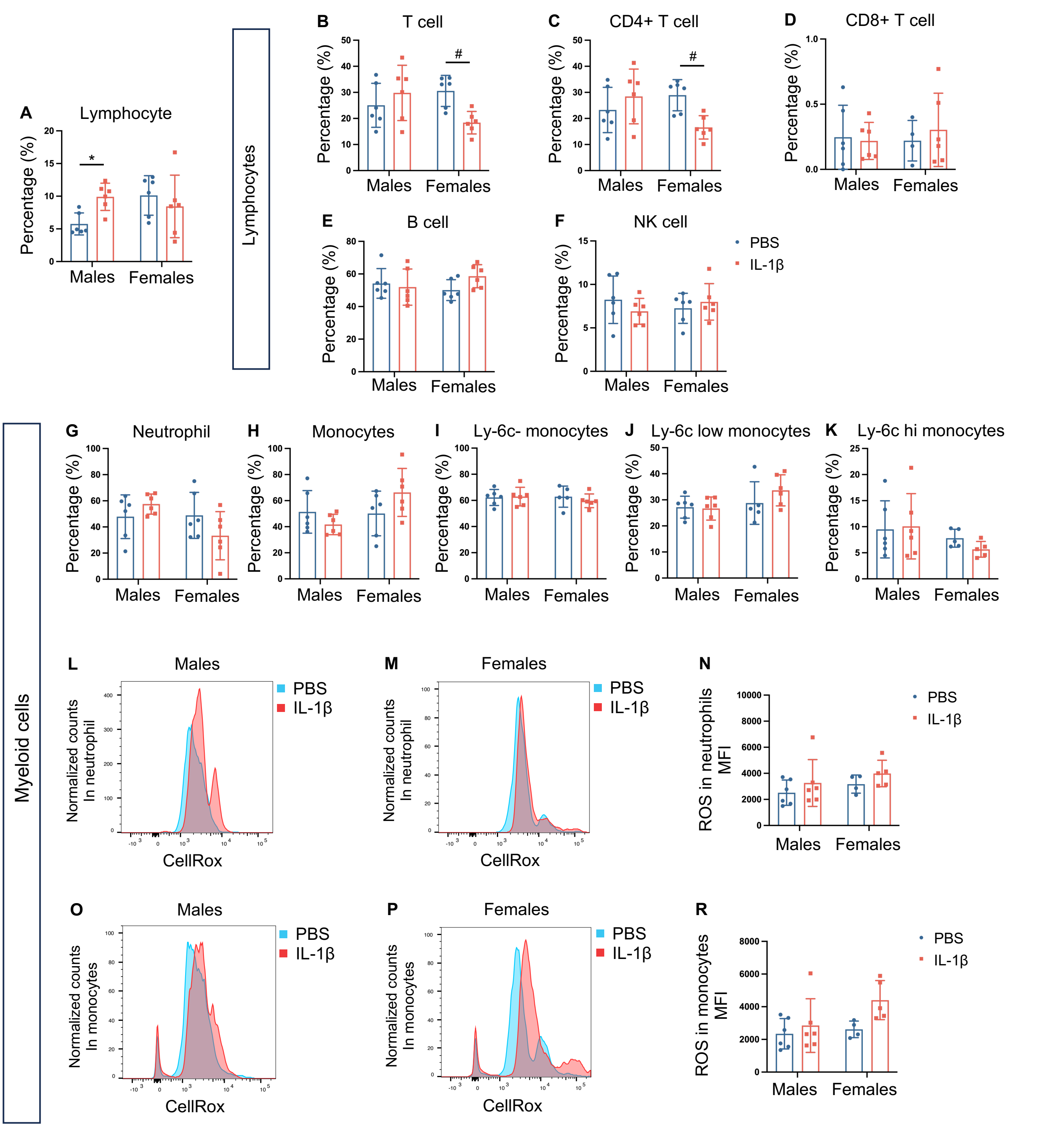

### Supplementary Figure 8

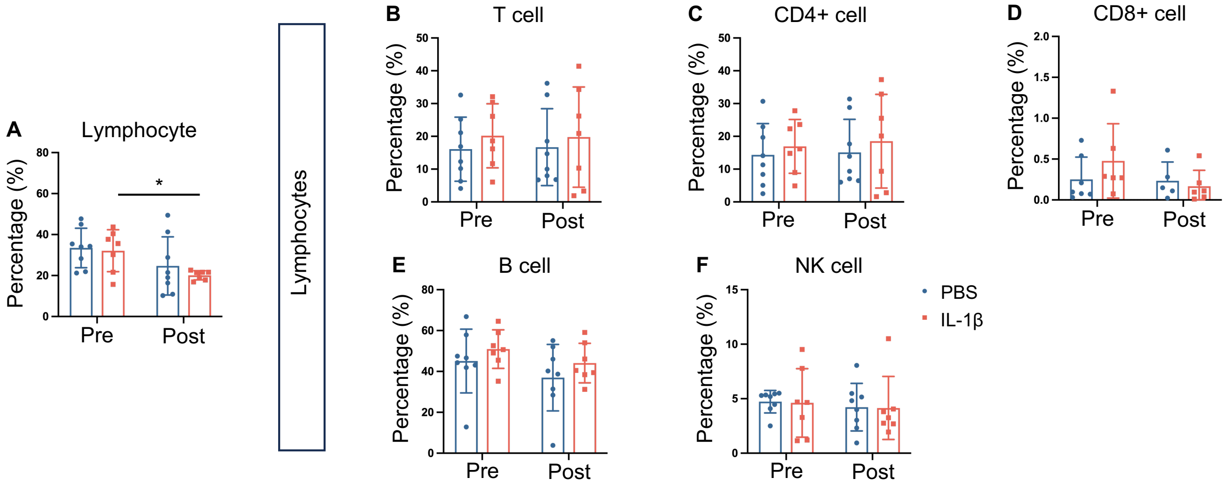
